# Design of proteins presenting discontinuous functional sites using deep learning

**DOI:** 10.1101/2020.11.29.402743

**Authors:** Doug Tischer, Sidney Lisanza, Jue Wang, Runze Dong, Ivan Anishchenko, Lukas F. Milles, Sergey Ovchinnikov, David Baker

## Abstract

An outstanding challenge in protein design is the design of binders against therapeutically relevant target proteins via scaffolding the discontinuous binding interfaces present in their often large and complex binding partners. There is currently no method for sampling through the almost unlimited number of possible protein structures for those capable of scaffolding a specified discontinuous functional site; instead, current approaches make the sampling problem tractable by restricting search to structures composed of pre-defined secondary structural elements. Such restriction of search has the disadvantage that considerable trial and error can be required to identify architectures capable of scaffolding an arbitrary discontinuous functional site, and only a tiny fraction of possible architectures can be explored. Here we build on recent advances in de novo protein design by deep network hallucination to develop a solution to this problem which eliminates the need to pre-specify the structure of the scaffolding in any way. We use the trRosetta residual neural network, which maps input sequences to predicted inter-residue distances and orientations, to compute a loss function which simultaneously rewards recapitulation of a desired structural motif and the ideality of the surrounding scaffold, and generate diverse structures harboring the desired binding interface by optimizing this loss function by gradient descent. We illustrate the power and versatility of the method by scaffolding binding sites from proteins involved in key signaling pathways with a wide range of secondary structure compositions and geometries. The method should be broadly useful for designing small stable proteins containing complex functional sites.

## Main Text

Many protein-protein interactions primarily involve a single contiguous structural element such as an alpha helix or an ordered loop. For example, the BCL2 family of anti-apoptotic proteins bind the BIM helix, botulinum toxin primarily interacts with a helical peptide from its cellular receptor, and many monoclonal antibodies recognize single contiguous peptides in their protein targets. In such cases, with modern protein design methods it is now straightforward to build proteins de novo that scaffold the relevant structural element, as has been done for the RSV-F epitope (*1*), anti-apoptosis inhibitors (*2*), and botulinum toxin (*3*), among others. When the binding elements span multiple structural elements, and are composed on one side primarily of alpha helices, as in the case of the IL2-IL2-receptor interactions, idealized helical bundles can be generated which place the relevant helices in the correct orientations (*4*). More general scaffolding of binding sites can be achieved using structure-blueprint-based approaches in which the binding interface segments are kept fixed and the connecting and terminal regions are built up using, for example, Rosetta fragment assembly (*5*), but achieving a viable protein architecture is challenging even if the locations of the secondary structure elements in the built-up region are specified. This is because the fragment assembly approach does not have an internal measure of global consistency of the placement of the various structural elements, so large numbers of backbones have to be built up and then subjected to CPU-intensive sequence design calculations to identify those for which plausible amino acid sequences likely to encode the structure can be designed.

To overcome this challenge, a method is needed to generate structures that contain the binding elements arrayed correctly in space that are scaffolded in a protein architecture that is likely to be designable -- that is, the lowest energy state of some amino acid sequence. We reasoned that the recently developed trRosetta residual neural net which maps protein sequences to residue-residue distance and orientation distributions (which can be readily transformed into protein 3D structures) could be adapted for this purpose (*6*). The trRosetta network, although trained entirely on native protein sequences and structures, predicts the structure of *de-novo-*designed proteins remarkably well. This made it possible recently to use the network to hallucinate completely new proteins without any constraint on the backbone geometry (*7*), and to design amino acid sequences likely to fold into a fully specified input backbone structure (*8*).

### Partially constrained protein hallucination with a composite loss function

To design proteins with a defined binding interface without having to specify overall structure, we explored the use of a loss function with two parts: a motif satisfaction loss, which captures the extent to which the sequence specifies the structure of the desired functional site, and a free hallucination loss, which captures the extent to which the sequence specifies a globally well-folded protein (Methods). Given a sequence and the coordinates of a specified functional motif, which can be composed of multiple discontinuous chain segments (Figure 1A, Methods), the trRosetta network predicts from the sequence C-β distance and orientation distributions between all pairs of residues (Figure 1B). The motif satisfaction loss is defined as the cross-entropy between the predicted distributions and those of the input structural motif, and rewards recapitulating the binding motif in the designed protein. The free hallucination loss is defined as the contrast (Kullback-Leibler (KL) divergence) between the predicted distributions and those from a sequence-agnostic background model, and is applied to the parts of the protein outside of the functional site. These two losses were previously used individually for fixed backbone sequence design (*8*) and *de novo* protein hallucination (*7*), respectively.

**Figure 1.**
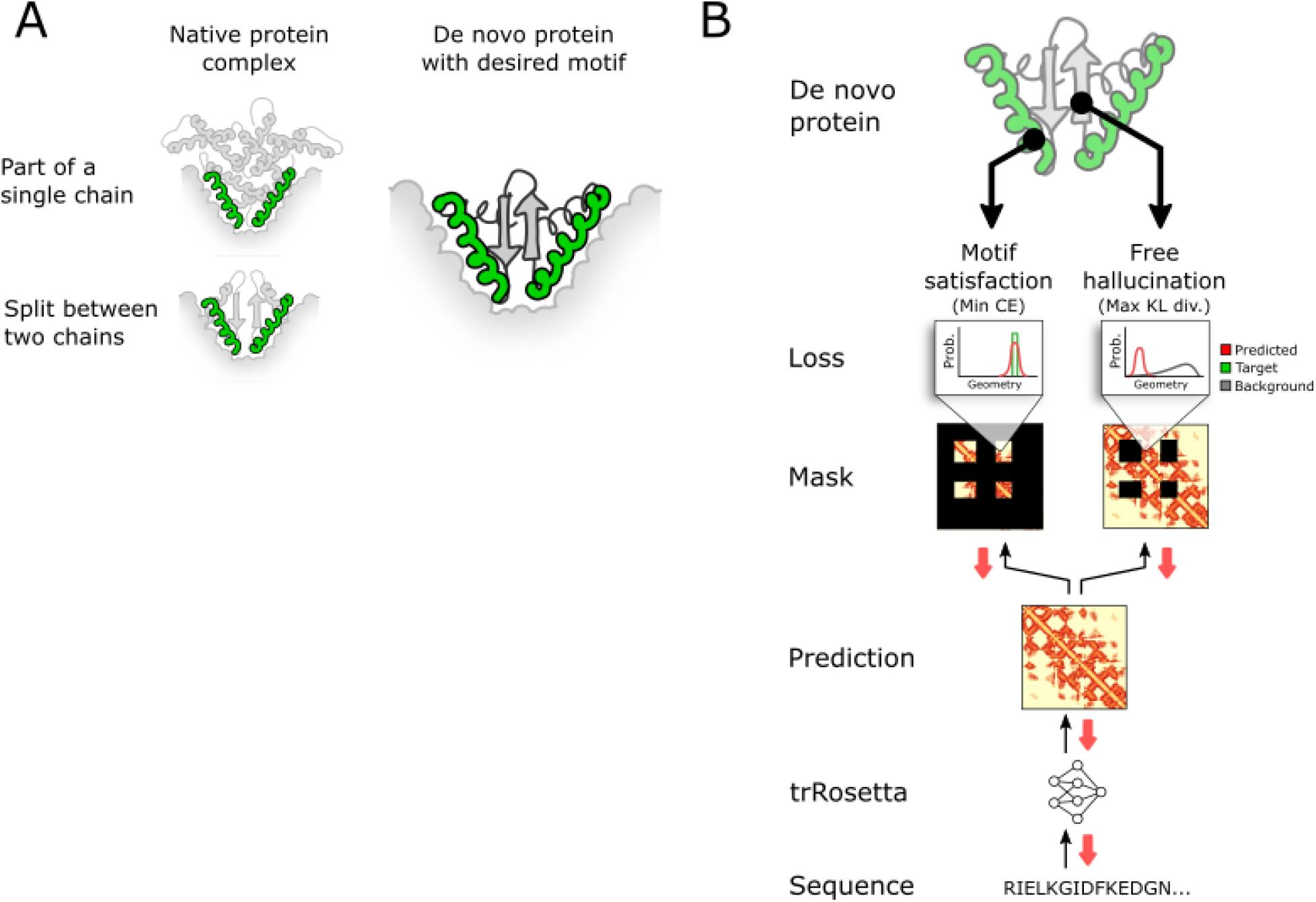
Incorporating multipartite geometric motifs into hallucinated proteins with a dual loss function. Protein binding motifs are important for a variety of biochemical functions, including protein complex assembly, substrate recognition and targeting therapeutic delivery. **(A)** Unfortunately, they can also be difficult to incorporate into a *de novo* protein because they can be part of a large, complex structure or split between two chains. Our technique seeks to build de novo scaffolds where the motif requirements are part of the design process from the beginning. **(B)** The forward pass in our network (black arrows) begins by feeding a random sequence to trRosetta, which predicts distributions of distances and angles between every residue. This prediction is masked and subject to the two mutually exclusive terms of the loss functions. The motif satisfaction term is the categorical cross entropy between the predicted distribution and the motif one-hot encoding and the free hallucination term is the negative KL divergence between the predicted distribution and a background distribution, which represents the average trRosetta prediction for a protein of the given length.The gradient with respect to the total loss is then calculated by backpropagation (red arrows) to the input sequence logits. The gradient is used to update the sequence and the process begins for the next step.

The evaluation of the motif satisfaction loss requires assigning residues in the protein sequence to functional site positions. We explored two approaches for carrying out this assignment. In the first approach, the sequence order of the functional site residues was preserved, and variable length sequence segments were placed before, between, and after blocks of residues assigned to the functional site. Independent optimization trajectories were then carried out, each initialized with stochastically chosen length for each inserted sequence segment from a user specified range (see Methods). In the second approach, the overall length of the sequence was fixed but the residues corresponding to the functional site were not prespecified; instead during optimization the functional sites were assigned to regions that best minimized the motif satisfaction loss, using a greedy search algorithm (see Methods).

We experimented with two approaches for optimizing the composite loss function (Figure 1B) starting from a randomly generated amino acid sequence. First, we explored a Monte Carlo sampling procedure in which starting at each iteration, a randomly selected amino acid substitution is made and the trRosetta network is used to generate predicted distances and orientations. The composite loss function is evaluated and the move accepted or rejected according to the standard Metropolis criterion. For proteins around 120 residues long, we found this approach converged in about 30,000 steps and took about 90 minutes on Nvidia GeForce RTX2080 GPUs.

Second, we evaluated gradient-based sequence optimization. Unlike the Monte Carlo approach, the gradient-based approach could design an entire multiple sequence alignment (MSA) because the gradient updates all sequence positions simultaneously, instead of mutating a single amino acid at time. We represented the MSA as a continuous random variable *Y_NxLxA_* (“input logits”) initialized randomly from Normal(0, 0.01), where *N* is the number of sequences in the MSA, *L* is the length of the protein and *A*=20 is the number of amino acids. On each iteration, we converted these input logits to amino acid probabilities by a softmax operation and one-hot-encoded them for input to trRosetta by either taking the most probable amino acid (i.e. argmax) or sampling from the amino-acid distribution (*9*) at each position in each sequence. To backpropagate the gradient to the continuous input logits through the discrete one-hot protein sequence, we employed a reparameterization trick (*8*, *9*), where gradients were passed through the one-hot sequence as if it had the softmax values of the input logits. Conceptually similar approaches are used in other works optimizing biological sequences (*10*, *11*). Over the course of a trajectory, we often decayed the learning rate according to a schedule, as it struck a balance between design quality and computational efficiency, but other optimization methods were superior in some cases (Methods). We found that gradient-based optimization with decay converged in about 200 steps for proteins about 120 residues long, taking approximately 5 minutes on our GPUs.

The output of optimization is a structure that scaffolds the binding interface and a sequence predicted to encode it. Because trRosetta does not have full atomic resolution (only the backbone is considered), we locally optimized the sequence and structure of each design in complex with the target structure using Rosetta full atom flexible backbone design, keeping the interacting residues inherited from the original binder fixed. This ensures tight complementary core packing, and improves shape complementarity with the target.

### Hallucinating scaffolds supporting functional sites

We tested the method by designing mimetics of a variety of naturally occuring proteins with structurally diverse binding interfaces (Figure 2, Table 1). We began by focusing on interfaces composed primarily of alpha-helices. We chose as representatives of such interfaces the complement cascade protein C3d which enhances immune responses to fused antigens (*12*), and the interleukin 4 (IL-4) cytokine which plays an important role in immune cell differentiation (*13*). In both cases, the method generated backbones which recapitulated the target motifs to within 1 Å Cα RMSD (Figure 2A-B). The generated designs have an additional 1-2 helices buttressing the interface; a hydrophobic core between the original helices and the added helices stabilizes the hallucinated folds. The C3d mimetics are much smaller (approximately 100 residues with 3-4 helices) than the native protein (307 AAs, 12 helices), a potential advantage for vaccine and other therapeutic applications (*4*, *14*).

**Table 1.** Natural proteins used for mimetic design.

| **Native protein** | **PDB ID** | **Chain** | **Constrained residues** | **Binding partner(s)** | **Reference** |
| --- | --- | --- | --- | --- | --- |
| IL-4 | 3BPL | A | 5-15, 78-92, 114-128 | IL4 receptor α-chain and cytokine receptor common γ-chain | (38) |
| Insulin | 6PXV | F | 10-19, 56-74 | Insulin receptor | (20) |
| C3d | 1GHQ | A | 104-126, 170-185 | Complement receptor 2 | (39) |
| GM-CSF | 4RS1 | A | 40-65, 97-120 | GMR-α | (23) |
| IL-10 | 1Y6K | A,B | 19-57, 133-155 | IL-10 receptor 1 | (17) |
| B7-2 | 1I85 | B | 28-32,40-43,85-89,95-100 | CTLA-4, CD28 | (22) |
| EF-hand | 1A75 | A | 88-97 | Ca<sup>2+</sup> | Unpublished |

**Figure 2.**
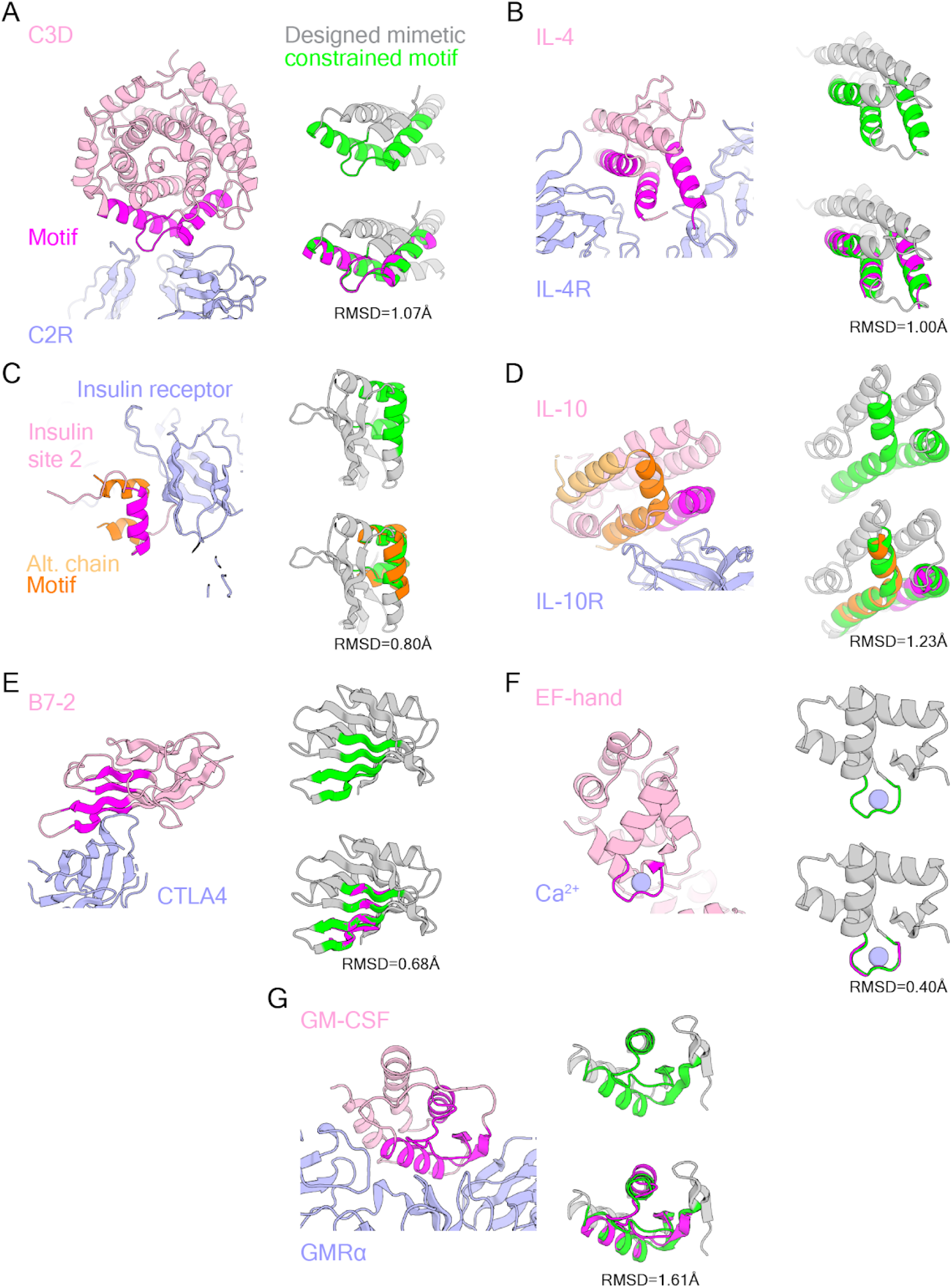
Design of scaffolds presenting discontinuous binding interface motifs of diverse natural ligands. Crystal structures of native binder/receptor complexes and examples of designed structures harboring the function site of interest. **(A and B)** Helical interfaces: C3d (PDB ID 1GHQ) **(A)** and IL-4 (PDB ID 3BPL) **(B)**. **(C and D)** Interfaces formed from different chains: insulin (PDB ID 6PXV) **(C)** and IL-10 (PDB ID 1Y6K) **(D)**. **(E)** The B7-2 interface formed primarily from β-strands (PDB ID 1I85). **(F)** The structured calcium binding EF-hand (PDB ID 1A75). **(G)** The GM-CSF (PDB ID 4RS1) interface is composed of a mixture of helices, sheets, and loops. In each panel, the native binding protein is colored light pink with the binding interface motif colored magenta (light orange/orange for the alternate chain) and the receptor or binding partner colored light blue. The designed protein is colored gray with the motif-mimicking region in green.

Since our hallucination approach does not rely on any information about the relative placement of the functional regions in the protein sequences from which they were derived, it can be readily applied to scaffold interfaces originally involving multiple chains. We chose as examples of such challenges the key blood sugar regulator insulin which is widely used in treating diabetes (*15*, *16*) and the anti-inflammatory cytokine interleukin 10 (IL-10) (*17*). Insulin is composed of two disulfide bonded chains derived from a larger precursor (pro-insulin) by proteolytic cleavage. The resulting unusual helical arrangement (Figure 2C, 3B) is challenging to recapitulate with traditional parametric helical bundle generation (*2*, *4*, *18*) or structure-blueprint-based methods in which the location of secondary structure elements in the sequence is specified, but not their relative orientation (*3*, *19*). Although insulin interacts with the insulin receptor through two binding sites (*20*), we focused on site 2 as it is formed primarily by alpha helices. The hallucination method generated mimics of this site (Figure 2C) that recapitulate the corresponding regions of insulin to 0.8 Å RMSD. IL10 is a domain-swapped dimer with two IL10R interacting domains each containing portions of the two chains in the dimer (*17*). We used the hallucination method to scaffold the binding site for a single receptor subunit in a single chain; the resulting scaffolds recapitulate the IL10 binding region to 1.23 Å RMSD (Figure 2D).

We next explored the scaffolding of functional loops with well-defined structures. We chose as an example the calcium binding EF-hand motif found in many naturally occurring calcium binding proteins and physiological processes (*21*). Unlike the other examples considered in this paper, the EF-hand is composed of a single contiguous chain segment, and the hallucination method readily generated a variety of scaffolds recapitulating the EF-hand motif to 0.4 Å RMSD. We tested a subset of these designs using Rosetta ab initio folding simulations to predict the structure from sequence. We found that the lowest energy states for the designs were very close to the hallucinated design models (Figure 4B,D). We carried out further *in silico* testing using explicit solvent Molecular Dynamics (MD), and found that many designs maintained calcium positioning inside the EF-hand loop in simulations longer than 300 nanoseconds (Figure 4A,C).

We next considered interfaces composed primarily of beta strands. We chose as an example the B7-2 protein (*22*), which is of considerable therapeutic relevance as it is a ligand for both the co-stimulatory T-cell surface protein CD28 and the inhibitory T-cell surface protein CTLA-4. We focused on scaffolding the B7-2 interface involved in CTLA-4 binding, which is composed of four beta strands (Figure 2E). Despite the greater complexity of folding beta sheet proteins, the hallucination method readily generated scaffolds recapitulating the binding interface to 0.68 Å RMSD. Rosetta ab initio folding simulations again provided independent in silico confirmation that the designed sequences encode the designed structures (Figure 4 F,H).

Finally, we investigated the scaffolding of more extensive interfaces composed by a mix of helices, strands, and loops. We chose as a case study the central hematopoietic growth factor and immune modulator granulocyte-macrophage colony-stimulating factor (GM-CSF) (*23*), which targets GMRα with an interface consisting of two strands and a helix. Despite the complexity of the interface, the hallucination method generated structures to 1.61 Å RMSD (Figure 2G).

### Structurally diverse proteins can scaffold the same motif

For each functional motif, the generated designs spanned a range of topologies (Figure 3). The IL-4 mimetics incorporate an additional helix to maintain the spatial positioning of the three binding helices, but there is considerable variation in the placement of the different helices in the sequence and structure (Figure 3A). The insulin mimetics solve the challenge of scaffolding helices from the two chains of insulin by positioning them within mixed α/β scaffolds with variable numbers of strands and helices varied; in all cases the network favored buttressing the binding helices with strands which were further stabilized by back helices (Figure 3B). In the case of B7-2 mimetics, which are required to scaffold four strands with strong curvature, solutions with quite different folds were obtained: some had all beta Ig-like folds similar to natural B7-2; others had alpha-beta ferredoxin-like folds (Figure 3C). The generated scaffolds were often quite different from any naturally occurring protein structure, with TM scores to the closest native protein in some cases below 0.6 (as expected, designs with smaller constrained regions generally had lower TM-scores, as trRosetta has more freedom to hallucinate in such cases).

**Figure 3.**
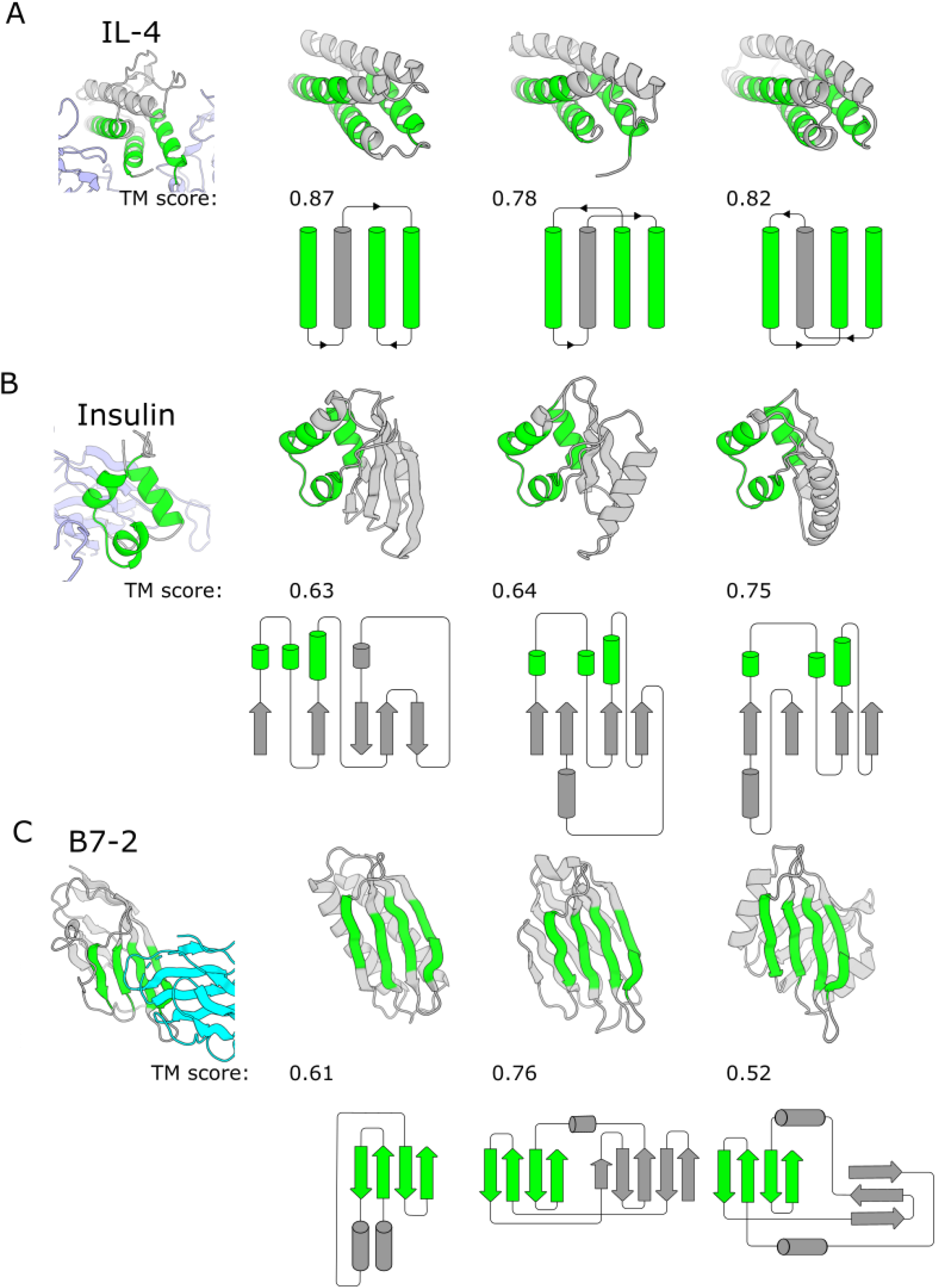
TrRosetta hallucinates diverse solutions to scaffolding the same functional motif. Three solutions are shown for **(A)** IL4, **(B)** insulin site 2, or **(C)** B7-2 binding motifs. The left image shows the native protein structure, with the binding motif colored green and the native binding partner colored blue. To the right of the native structure are ribbon diagrams of three trRosetta solutions, with the region constrained by the motif geometry colored green. The reported TM-score is the highest score to any protein in the PDB, indicating that trRosetta didn’t simply generate the structure of the native protein or any other protein in the PDB (TM-scores are higher when the geometric motif is larger, because a larger part of the hallucinated protein is constrained). Topology diagrams show that although the tertiary structure of design solutions can be similar, their secondary structure elements are connected in very diverse ways; the method does not suffer from mode collapse, which can occur with other generative techniques such as GANs.

**Figure 4.**
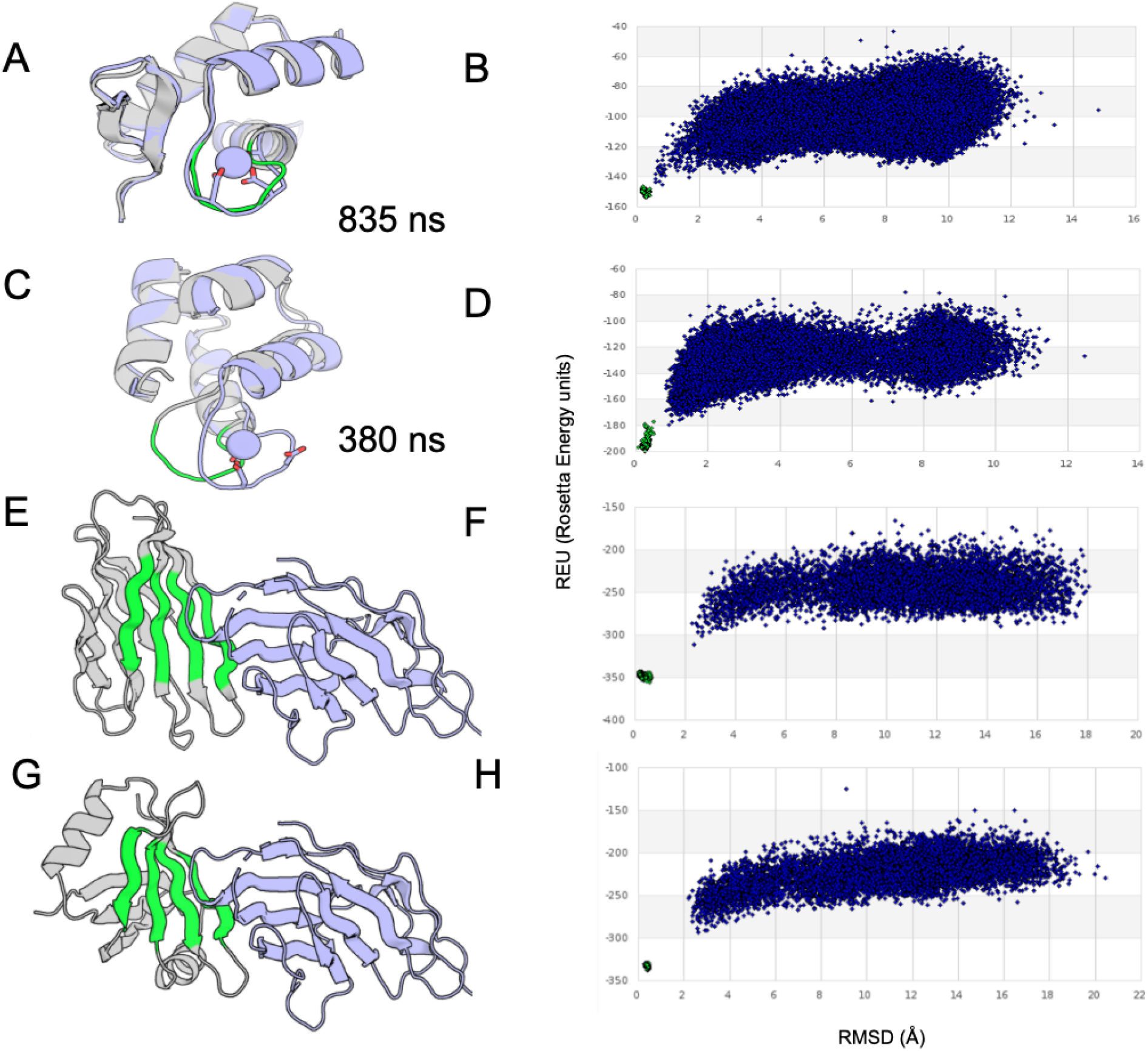
In silico validation of hallucinated proteins. **(A,C,E,G)** Hallucinated design models presenting the EF-hand loop in grey **(A,C)** or the CTLA-4 binding B7-2 mimetics **(E,G)**, with the scaffolded region shown in green. **(B,D,F,H)** Energy landscapes for the designed sequences for the structures shown in **(A,C,E,G)** mapped using Rosetta de novo folding simulations (y axis energy, x axis distance from design model). In each case, the lowest energy structure is very close to the design model, providing independent in silico validation of the folding of the sequence to the designed structure. **(A,C)** Starting and final snapshot of 380 and 835ns MD simulations with calcium for EF hand designs in grey and cyan respectively.

## Conclusions

We have described a deep-learning-based method to generate protein scaffolds containing a predefined functional motif that spans multiple discontinuous structural elements. The method requires no inputs other than a desired sequence length and the structure of the functional motif, thus offering much greater flexibility over existing methods that restrict the secondary structure of the motif, the topology of the scaffold, or both. Full confirmation of the accuracy and robustness of our approach will require experimental characterization of folding and binding, but the designed sequences are predicted to fold to the designed structures in *ab initio* forward-folding simulations, a stringent *in silico* benchmark.

Our approach further illustrates the broad utility of “inverted models” which optimize an input sequence with respect to a loss function, and have previously been used to design DNA sequences (*10*), guide directed evolution (*24*), and design proteins using trRosetta (*7*, *8*, *11*). An alternative deep-learning paradigm generates proteins directly in the forward pass of a generative adversarial network (GAN) or variational autoencoder (VAE) (*25*). Although GANs and VAEs have been used to design functional sequences (*26*–*28*) or biophysically plausible structures (*29*, *30*), none have so far been used to create new proteins with functional elements. Furthermore, VAEs have only been successfully trained on families of related proteins (*30*), whereas trRosetta has been trained on, and thus can in principle generalize from, all known protein structures. Finally, existing GANs require retraining to accommodate varying sequence lengths (*29*). By contrast, with an inverted model, loss functions and design hyperparameters such as sequence length can be varied without retraining.

Despite its broad scope and ability to rapidly search through structure space for solutions, our approach is currently limited by the accuracy of trRosetta structure predictions, about 2 Å RMSD on average (*6*, *8*). To overcome this limitation, we currently refine the sequence and structure of the designs generated by the network using Rosetta all-atom flexible-backbone design calculations. In essence, we use deep network hallucination to solve the open-ended architecture and fold design sampling and design problems, and Rosetta for the higher resolution details. As progress in deep-learning-based protein structure prediction continues, we expect it to become possible to directly generate more and more accurate functional designs using our dual-loss hallucination approach without need for further refinement.

## Methods

### Sequence representation

For structure prediction, the input to trRosetta is a one-hot-encoded multiple-sequence alignment (MSA) in the form of an N x L x A tensor, where L is the length of the first sequence in the alignment and the protein whose structure is being predicted, N is the number of aligned sequences, and A=21 is the number of amino acids plus gap character (although we do not use gaps during design). TrRosetta accurately predicts the structure of *de novo* proteins even when only one sequence is used as input (N=1), so previous work on unconstrained protein design optimized a single sequence (*7*). When designing proteins to recapitulate structural constraints derived from natural backbones, however, better accuracy was obtained when the loss was optimized over an entire MSA (*8*), probably due to the increased degrees of freedom. Therefore, in this work, we optimized an MSA of N=1000 sequences to generate each scaffold, with an 80% dropout on the input features to TrRosetta (including 2D tiled sequence, conservation and coevolution) to avoid overfitting.

### Loss function

Our composite loss function (Figure 1B) scores how well a given sequence satisfies our dual design constraints. That is, does the given sequence strongly encode a backbone geometry and does that backbone geometry include the binding site motif. It contains a motif satisfaction (MS) loss and a free hallucination (FH) loss.

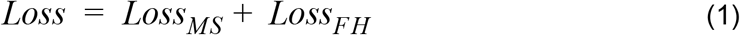

The motif satisfaction loss is the cross entropy of the network predictions at the coordinate values (*y*^0^, *y* ∈{*d*, ω, θ, φ, θ^*T*^, φ^*T*^}) derived from the target motif, averaged over all residue pairs in the masked regions (*m*).

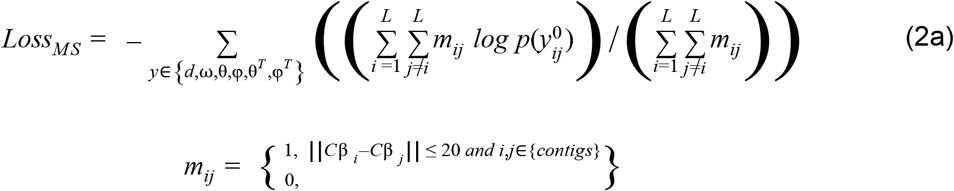

The free hallucination loss is the KL divergence between the network predictions (*y*) and a background distribution (*b*) (*7*).

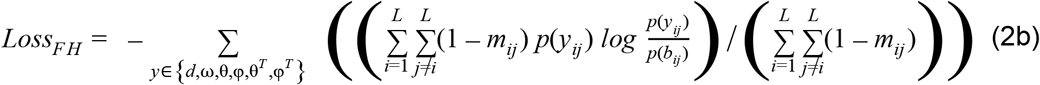

We explored several optimization procedures to find sequences that minimized this loss.

### Optimization methods

First, we explored a Monte Carlo sampling procedure similar to that employed in our de novo hallucination study. Starting from a random sequence, single mutations were proposed and the loss function evaluated. The mutation was either accepted or rejected according to the standard Metropolis criterion. We found that this approach converged in roughly 30,000 steps for a 120 amino acid protein with a 50 residue motif, which took about 90 minutes on our GPUs. Although slow, this approach has the advantage that mutations can include insertions and deletions, which allows for the lengths of loops to be easily optimized.

Second, we evaluated a gradient based sequence optimization similar to that used in our fixed backbone sequence design study (*8*). Starting with randomly initialized input logits (*Y*; *N(0, 0.01)*), the gradient of the loss function with respect to the upstream logits is computed by backpropagation through the network. We explored applying the gradient to the input logits by normalizing it and applying a constant learning rate

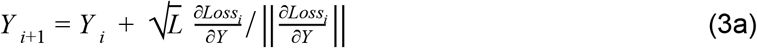

(3a)or by scaling the learning rate according to a predefined schedule (gradient decay).

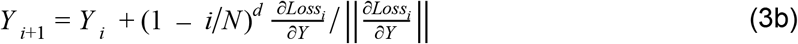

where *L* is the length of the protein, *i* is the optimization step, *N* is the total number of optimization steps and *d* is the decay rate.

We found that both methods converged to roughly the same range of loss values in 100 iterations for a 120 residue protein, taking about 3 minutes on our GPUs. We defaulted to using the gradient decay method for the structures in this paper, unless noted otherwise.

### Hardware

Tensorflow code for trRosetta was run on Nvidia GeForce RTX2080 GPUs.

### Motif placement algorithms

We applied one of two motif placement algorithms; one which we specify a range of lengths for the regions to be hallucinated between motifs before optimization and the other which determines motif placement during optimization via a greedy search. We use “contig” to refer to a contiguous portion of the designed sequence that is subject to the motif-satisfaction loss.

#### Pre-specified motif placement

The first algorithm takes the input structure and specifies multiple regions to replace with a new hallucinated region of a given length range. A specific value within each range is chosen randomly before starting optimization. The free hallucination loss is applied to these regions, while the motif satisfaction loss is applied to the rest of the structure. The residue positions being hallucinated or constrained as motifs stay fixed throughout each optimization trajectory, but many trajectories are run with different sampled lengths of the hallucinated regions. This approach allows for greater control over the tertiary structure of designs. It also allows users to shorten connections between contigs, limiting the search space, while lengthening the connections in other regions, biasing the network to do a more exhaustive conformational search in these regions.

#### Motif placement during optimization via greedy search

Other times it may not be obvious *a priori* how the contiguous segments of the motif (contigs) should be joined, especially if the contigs are small or far apart. As an alternative to running many trajectories that sample different lengths for these intervening gaps, we used a greedy search algorithm to determine where to apply the motif satisfaction loss.

The idea behind the algorithm is to apply the motif satisfaction loss to regions in the predicted geometry that already closely match the target geometry at each step of the optimization. Initially, the motif satisfaction loss is applied somewhat randomly but then eventually settles on one region once it starts to minimize towards the desired geometry. From there, the process becomes self-reinforcing as the motif satisfaction loss will be applied to regions that are already similar to the desired geometry.

We focused the search on regions in the predicted geometry that minimize the cross entropy of the *inter*-contig geometry because we were primarily focused on how to space the contigs relative to each other. Regions where the contigs should be placed (and the corresponding motif satisfaction loss applied) was determined in a stepwise manner.

In the first step, we placed two contigs simultaneously because we searched the predicted geometry with the inter-contig geometry (the one-hot encoding of the 6D transform between all pairs of residues between two contigs). We convolved the inter-contig geometry over the negative log of the network predictions, thus calculating the cross entropy for all possible placements of a contig pair. We required contigs to remain in a user defined order. Positions that would result in the contigs overlapping with each other or prevent the placement of the remaining contigs were scored as positive infinity. This process was repeated for all contig pairs, after which the contig pair with the lowest cross entropy was fixed in place.

In the second step, contigs were placed one at a time until none were left. A new contig was placed such that it minimized the inter-contig cross entropy with the already-placed contigs.

Because greedy searches can miss global optimums, we added the top 3 scoring results at each step to a search tree, yielding a collection of possible contig placements. The contig placement with the lowest motif satisfaction loss was then used to mask the network predictions (*m* in equation 2a), dividing it into parts subjected to the motif satisfaction or free hallucination loss.

### Designing mimetics of natural protein ligands

We selected binding motifs from protein interfaces that contained mostly secondary structural elements and were biomedically relevant. First, we identified interface residues as those within 5 Å of the binding partner. We made the binding motifs by manually clustering these binding residues (and any intervening residues) into several contiguous elements. In some cases, we also manually added other contiguous elements from the native structure to buttress the interface geometry. Table 1 lists the mimetic design targets, their PDB accessions, the residue numbers of constrained regions, and references.

### Rosetta sequence design and *ab initio* folding

Briefly, we used the Rosetta FastDesign mover with layer design and/or fragment based PSSMs to constrain amino acid choices in the protein. We also constrained residues that formed key interaction in the native protein structure (both to its binding partner and between regions of the binding motif) to only repack. We added harmonic potential restraints to these key residues to ensure they didn’t move during relaxation. The complete scripts used for sequence design will be released with the full version of this paper.

After the sequences were designed, we used Rosetta *ab initio* folding to test the plausibility of the protein sequence to encode the desired structure. In some cases we prevented fragment insertion in motifs that had large loops as it was hard to find good quality fragments for those regions, even though we knew that geometry was possible from the native protein structure. The complete scripts for ab initio folding will be released with the full version of this paper.

### Molecular dynamics simulations

We equilibrated our model structures in explicit water using the NAMD (NAMD3alpha6) (*31*) molecular dynamics package with initial runfiles and setups created by VMD (*32*) plugin QwikMD (*33*). We used the CHARMM36 force field (*34*) and TIP3 water model (*35*) in all simulations. We centered structures in a water box at least 15 Angstrom larger than the protein’s longest dimension, NaCl was added to 150 mM. Minimization (2000 steps), then Annealing (0.29 ns, temperature rise 60 K to 300 K, 1 atm pressure, protein backbone restrained), then equilibration (1 ns, temperature 300 K, 1 atm pressure, protein backbone restrained), then we performed MD simulation (temperature 300 K, 1 atm pressure, no restraints) in the NpT ensemble. We ran the final MD equilibration simulations for at least 300 ns, in which an N- or C-terminal amino acid backbone far from the Ca-binding loop of each design was restraint harmonically to keep the design centered in the water box.

All simulation parameters were: a distance cut-off of 12.0 Å was applied to short-range, non-bonded interactions, and 10.0 Å for the smoothing functions. Long-range electrostatic interactions were treated using the particle-mesh Ewald (*36*) method. The pressure was maintained at 1 atm using Nosé-Hoover Langevin piston (*37*). The equations of motion were integrated using the reversible reference system propagator algorithm (r-RESPA) multiple time step scheme to update the short-range interactions every 1 steps and long-range electrostatics interactions every 2 steps. The time step of integration was 2 fs for all simulations. The temperature was maintained at 300 K using Langevin dynamics.

## Acknowledgements

We would like to thank Luki Goldschmidt for running and maintaining the computational resource in the IPD. Chris Norn and Justas Dauparas were helpful for general discussions about trRosetta and its capabilities.

## Funding Sources

D.T. is supported by The Open Philanthropy Project Improving Protein Design Fund. J.W. is supported by a postdoctoral fellowship from the Washington Research Foundation. S.L. is supported by Amgen. L.F.M. is supported by a Human Frontier Science Program Cross Disciplinary Fellowship (LT000395/2020-C) and an EMBO Non-Stipendiary Fellowship (ALTF 1047-2019). I.A. is supported by the National Institute of Allergy and Infectious Diseases (NIAID, Federal Contract HHSN272201700059C). S.O. supported by NIH grant DP5OD026389. D.B. is supported by the Howard Hughes Medical Institute.

## Notes

### Competing Interest Statement

The authors have declared no competing interest.

